# Precision Breathing: Asthma Phenotyping via Machine Learning

**DOI:** 10.1101/2025.04.24.650383

**Authors:** Nathan Kodjo Mintah Churcher, Antra Ganguly, Nana Kwame Ayisi-Boateng, Ernest Adankwah, Michael Kofi Ansah, Richard Odame Phillips, Shalini Prasad

## Abstract

Asthma is a complex condition characterized by chronic airway inflammation, with varying severity, symptoms, triggers, and treatment responses. Traditional classification relies on clinical attributes, but the growing understanding of asthma’s heterogeneity highlights the need for phenotyping. Effective management requires regular monitoring, medication, and prevention of exacerbations, but current diagnostic methods face challenges such as the lack of definitive tests and reliance on subjective measures. Implementing precision medicine, especially for severe cases, necessitates identifying measurable markers in biofluids. This study explores machine learning methods to identify biomarkers differentiating various asthma phenotypic states. We measured inflammatory markers in both plasma and saliva samples and used machine learning algorithms to determine their efficacy in reflecting airway inflammation. Our findings indicate that saliva markers provide a more accurate representation of localized inflammation compared to plasma markers, which reflect a systemic response. Using MRMR (Minimum Redundancy Maximum Relevance) ranking, we enhanced model efficacy. The K-Nearest Neighbor (KNN) classifier achieved 75% accuracy with the first 12 saliva markers, while the Random Forest (RF) classifier performed best for plasma models, though with lower accuracy. Our results suggest machine learning can effectively identify key markers for asthma phenotyping, aiding personalized treatment strategies. Customizable point-of-care devices could validate these models and improve their accuracy, advancing asthma treatment and management.

## Introduction

Asthma is a complex and heterogeneous condition of the lungs, usually associated with chronic inflammation of the airways that can vary in severity, symptoms, triggers, and response to treatment(1). Traditionally, classification of asthma has been based on clinical attributes such as the symptoms observed and response to treatment. However, with the growing understanding of the heterogeneity of asthma as a spectrum with varying underlying causes, asthma phenotyping has been on the horizon. Asthma management requires regular monitoring, medication, and prevention of exacerbations(2,3). However, there are many challenges and limitations in the current methods of asthma diagnosis, classification, and prediction. Current existing criteria are based on subjective and variable measures(4). Implementation of precision medicine in asthma management, especially in the case of severe or uncontrolled asthma, would necessitate the identification and study of measurable markers in biofluids.

The efficacy of treatment for asthma depends on the identification of the type of inflammation present in the patient due to the disease’s varied nature(5). Asthma inflammation is usually classified by the T-helper cells and chemical messengers involved. Th2 inflammation, which is more common, involves cytokines that attract eosinophils and is often linked to allergic asthma (6,7). On the other hand, Th1 inflammation is associated with neutrophilic asthma, which typically does not respond well to standard treatments(6,7). Chemokines and cytokines play crucial roles in immune responses and inflammation, particularly in asthma(8). They mediate communication between the cells in the immune system and as such have a direct effect on the inflammation process. The role of some cytokines has been established in the pathogenesis of asthma and due to the involvement of cytokines in the modulation of asthma, can be an excellent target for asthma phenotyping(6).

CCL3 (MIP-1α) and IP-10 (CXCL10) both attract immune cells to inflammation sites, with CCL3 recruiting monocytes, macrophages, and T cells, and IP-10 attracting Th1 lymphocytes(9–11). Similarly, IL-8 and G-CSF are involved in neutrophil recruitment, with IL-8 guiding neutrophils to lung tissue and G-CSF enhancing their production(12–15). IL-4 and IL-1α both promote eosinophil recruitment, contributing to airway inflammation and remodeling(16–19). IL-12 and IFN-γ are Th1 cytokines that can inhibit allergic airway inflammation, with IFN-γ potentially protecting against Th2-mediated asthma(20–22). TNF-α and CD40Ligand both promote inflammation by recruiting neutrophils and eosinophils and increasing pro-inflammatory cytokines(23,24). IL-2’s role in asthma is less clear, but elevated levels are noted in asthmatic patients(25). IL-6 acts as a pro-inflammatory cytokine linked to various asthma severities, while IL-10 suppresses airway inflammation but may hinder infection response(26,27). IL-17/IL-17A is associated with non-eosinophilic asthma and increased airway hyperresponsiveness(28). Additionally, Procalcitonin (PCT) is a marker for bacterial infections and can help determine if severe asthma cases are precipitated by bacterial lower respiratory tract infections, guiding antibiotic treatment to reduce hospitalization times and antibiotic use(29). TRAIL, a protein involved in programmed cell death, balances pro and anti-inflammatory cytokines, recruits eosinophils, and may induce apoptosis in airway epithelial cells, making it a potential therapeutic target for asthma management(9).

In this study, the selected cytokines were screened in blood plasma and saliva samples obtained different groups: Healthy Non-Smokers (HNS), Healthy Smokers (HS), Diseased Stable individuals (DS) clinically diagnosed with asthma but controlled without any recent flare-ups), and Diseased Acute individuals (DA) clinically diagnosed with asthma with a recent flare up or respiratory infection. Leveraging the cytokines measured in blood and saliva, a machine learning approach was adopted to identify a combination of the cytokines to assess the disease activity and further aid phenotyping of patients with asthma.

## Materials and methods

### Subject recruitment

This study recruited individuals aged 18 to 65 years from the outpatient unit of the university hospital at Kwame Nkrumah University of Science and Technology (KNUST). Eligibility was determined by trained medical personnel following predefined criteria. Participants were selected using convenience sampling, either through word-of-mouth or via flyers posted in the hospital’s outpatient wing. A total of 60 subjects were enrolled in the study, categorized as follows: 10 HNS, 10 HS, 20 DS, and 20 DA.

### Ethics statement

Studies were approved by the Institutional Review Board (IRB) at University of Texas at Dallas (UTD 21-466) and the Committee on Human Research, Publication and Ethics at KNUST (CHRPE/AP/519/21). Recruitment began on 12/06/2021 and ended on 03/06/2022. Written informed consent was obtained from all subjects as approved by the KNUST Committee on Human Research, Publication and Ethics.

### Collection procedures

#### Blood collection and processing

Approximately 10mL of venous blood was collected from each participant by a trained phlebotomist into EDTA tubes. All participants provided written informed consent for the collection of their blood samples. Samples were transported to Kumasi Centre for Collaborative Research, Kwame Nkrumah University of Science and Technology in Tropical Medicine (KCCR) for processing. Blood plasma was harvested by centrifuging at 2000 xg for 15 minutes at 4°C. The isolated plasma was transferred into sterile 1.5mL cyrovial tubes and stored in a −80°C freezer until they were transported to UT Dallas for biomarker quantification. Freeze-thaw cycles were avoided to preserve sample quality. All the samples were collected between the hours of 1pm and 6pm.

#### Saliva collection and processing

Approximately 2mL of saliva was collected by allowing saliva to drool into the vial after the mouthpiece of the saliva kit is inserted into the mouth of the participant. The collected samples were frozen in a −80°C freezer at KCCR until transported frozen to UT Dallas for biomarker quantification. Freeze-thaw cycles were avoided to preserve sample quality. Saliva samples were collected at the same time as blood samples. All the samples were collected between the hours of 1pm and 6pm.

#### Biomarker quantification

The concentrations of cytokines (G-CSF, IL-1α, IL-4, IL-8, IL-12, IFN-γ, TNF-α, IL-6, IL-10, IL- 17/IL-17A), chemokines (CCL3, IP-10), and other inflammatory markers (CD40 Ligand, PCT, TRAIL) in blood plasma and saliva were quantified using Luminex xMAP technology at UT Southwestern Medical Center. Luminex assays allow for the simultaneous detection of multiple analytes using color-coded microspheres conjugated with specific antibodies, providing high sensitivity and efficiency in biomarker analysis.

Prior to analysis, frozen plasma and saliva samples were thawed, centrifuged at 2000 × g for 5 minutes at 4°C, and the supernatant was extracted for quantification. Assays were performed according to the manufacturer’s protocol, ensuring reproducibility and accuracy in detection.

### Statistical analysis and Machine Learning

Graphpad Prism® was used to perform statistical analysis and data visualization. Google’s CoLab platform (quad core Intel Xeon processor, Tesla V100-SXM2-16GB GPU) was used for building machine learning models and visualizing the results. In-built library models in python were used for the respective classifiers used in assessing the utility of the markers to stratifying the 4 different groups, with bootstrapping was done randomly with replacement. The GridSearch from the sklearn library(30) with cross-validation was used to tune the different classifiers tested to achieve the maximum accuracy possible. Feature selection was decided based on the minimum Redundancy - Maximum Relevance (mRMR) ranking on the feature importance score calculated intrinsically by finding the smallest relevant subset of features for a given task (31). Data visualization was done using Matplotlib and Seaborn functions.

## Results and Discussion

The naïve expression of the cytokine panel was assessed in both plasma and saliva to understand the relationship of cytokine expression in both a systemic and relatively localized expression in the case of asthma. Among the biomarkers measured in both plasma and saliva, some values were either below the limit of detection (OOO) or required extrapolation. Table 1 presents the number and percentage of such measurements for both biofluids. As expected, more than 50% of plasma markers were either undetectable or extrapolated, likely due to their lower concentrations in plasma compared to saliva in assessing respiratory conditions such as asthma. To maintain data integrity, markers with more than 55% of values classified as OOO or extrapolated were excluded from the dataset, as indicated in Tables 1 and 2. This exclusion criterion ensures the reliability of the analysis by focusing only on markers with detectable and quantifiable levels.

**Table 1.** Out of Range and Extrapolated rate of the marker panel in saliva and plasma.

|  | Plasma |  | Saliva |  |
| --- | --- | --- | --- | --- |
| Inflammatory Marker | OOOR<br>n (% of total) | Extrapolated<br>n (% of total) | OOOR<br>n (% of total) | Extrapolated<br>n (% of total) |
| CCL3 | 78 (92.86%) | 1 (1.19%) | 0 (0.00%) | 0 (0.00%) |
| IP-10 | 0 (0.00%) | 0 (0.00%) | 1 (2.50%) | 7 (17.50%) |
| G-CSF | 66 (78.57%) | 14 (16.67%) | 0 (0.00%) | 0 (0.00%) |
| IL-1ALPHA | 83 (98.81%) | 1 (1.19%) | 0 (0.00%) | 2 (5.00%) |
| IL-1ra | 0 (0.00%) | 0 (0.00%) | 20 (50.00%) | 20 (50.00%) |
| IL-4 | 53 (63.10%) | 0 (0.00%) | 0 (0.00%) | 0 (0.00%) |
| **IL-8** | 0 (0.00%) | 30 (35.71%) | 0 (0.00%) | 14 (35.00%) |
| **IL-12p70** | 66 (78.57%) | 17 (20.24%) | 0 (0.00%) | 15 (37.50%) |
| **IL-31** | 83 (98.81%) | 1 (1.19%) | 16 (40.00%) | 9 (22.50%) |
| **TNF-Alpha** | 0 (0.00%) | 44 (52.38%) | 0 (0.00%) | 1 (2.50%) |
| **CD40Ligand** | 0 (0.00%) | 11 (13.10%) | 0 (0.00%) | 3 (7.50%) |
| **D-dimer** | 0 (0.00%) | 2 (2.38%) | 1 (2.50%) | 3 (7.50%) |
| **IFN-Gamma** | 13 (15.48%) | 68 (80.95%) | 0 (0.00%) | 4 (10.00%) |
| **IL-1Beta** | 79 (94.05%) | 5 (5.95%) | 1 (2.50%) | 0 (0.00%) |
| **IL-2** | 61 (72.62%) | 23 (27.38%) | 0 (0.00%) | 1 (2.50%) |
| **IL-6** | 23 (27.38%) | 49 (58.33%) | 0 (0.00%) | 0 (0.00%) |
| **IL-10** | 64 (76.19%) | 4 (4.76%) | 0 (0.00%) | 0 (0.00%) |
| **IL-17/IL-17A** | 64 (76.19%) | 20 (23.81%) | 0 (0.00%) | 10 (25.00%) |
| **Procalcitonin** | 0 (0.00%) | 1 (1.19%) | 0 (0.00%) | 0 (0.00%) |
| **TRAIL** | 44 (52.38%) | 7 (8.33%) | 0 (0.00%) | 0 (0.00%) |

**Table 2.** Marker panel and summary statistics for plasma and saliva.

| Inflammatory Marker | Mean ± SD pg/mL | Median (Range) pg/mL |
| --- | --- | --- |
| **Plasma** |  |  |
| **IP-10** | 66.72 ± 92.54 | 39.57(13.94-447.38) |
| **IL-1ra** | 307.28 ± 180.72 | 259.93(54.15-1202.47) |
| **IL-8** | 3.56 ± 2.96 | 2.83(0.04-14.71) |
| **TNF-α** | 4.91 ± 3.72 | 4.1(1.36-23.11) |
| **CD40Ligand** | 317.02 ± 295.52 | 208.94(30.12-1835.18) |
| **D-dimer** | 1812895.5 ± 2463555.78 | 1053700(139269.5-14143000) |
| **Procalcitonin** | 40.85 ± 48.85 | 26.25(4.21-318.81) |
| **Saliva** |  |  |
| **CCL3** | 403.33 ± 85.77 | 377.17(269.33-627.89) |
| **IP-10** | 448.18 ± 667.50 | 238.56(16.47-3697.22) |
| **G-CSF** | 226.3 ± 295.22 | 124.86(18.42-1352.30) |
| **IL-1α** | 767.47 ± 906.14 | 454.08(49.08-4621.87) |
| **IL-4** | 158.89 ± 197.18 | 102.84(72.92-1123.70) |
| **IL-8** | 1818.23 ± 1753.68 | 1307.79(223-8154.49) |
| **IL-12** | 108.84 ± 33.81 | 105.49(51.66-182.22) |
| **IL-31** | 242.95 ± 388.09 | 82.98(3.37-1573.7) |
| **TNF-α** | 45.42 ± 80.75 | 15.35(3.31-366.03) |
| **CD40Ligand** | 501.83 ± 808.82 | 248.96(90.63-4286.37) |
| **D-dimer** | 2727934.44 ± 2712800.97 | 1980000(4309.36-10600000) |
| **IL-1β** | 811.39 ± 850.45 | 537.99(70.16-4314.65) |
| **IL-2** | 78.77 ± 47.61 | 69.26(19.55-215.72) |
| **IL-6** | 14.67 ± 16.39 | 9.55(2.81-101.6) |
| **IL-10** | 50.43 ± 23.45 | 42.05(9.95-128.2) |
| **IL-17/IL-17A** | 62.38 ± 162.69 | 12.73(5.35-978.25) |
| **Procalcitonin** | 22.81 ± 10.99 | 20.04(12.42-59.77) |
| **TRAIL** | 1074.09 ± 879.38 | 823.72(163.99-4470.64) |

*IP-10* Interferon-gamma inducible protein 10, *IL* interleukin, *TNF-α* tumor necrosis factor-alpha, *CCL3* Chemokine (C-C motif) ligand 3, *G-CSF* granulocyte colony-stimulating factor

Descriptive statistics for the different markers in both plasma and saliva have been presented in Table 2. In accordance with literature, we observed relatively higher concentrations (both mean and median, except for procalcitonin) of the inflammatory markers in saliva compared to plasma(32–35). The higher concentrations of inflammatory markers in saliva compared to plasma suggests that saliva may be a more sensitive indicator of localized inflammation in the airways. This could lead to the development of non-invasive diagnostic tools using saliva samples to monitor asthma severity and response to treatment.

The measurements were taken for all four groups (Healthy Non-Smoker, Healthy Smoker, Diseased Stable, Diseased Acute). While for most markers (see Supplementary Information), there was an increased expression in the disease group compared to the healthy cohort, there was no statistically significant difference observed between the groups. This lack of statistical significance highlights the complexity of cytokine expression patterns and the need for a more sophisticated analysis. Employing machine learning techniques to analyze multiplexed cytokine data can help identify patterns and interdependencies that are not apparent through traditional statistical methods. Machine learning can help identify subtle patterns and interactions among the cytokines providing a more comprehensive understanding of the inflammatory responses in asthma.

To analyze the non-linear relationships between the biomarker expression for both biofluids, saliva and plasma, we built several supervised Machine Learning (ML) models. Since two biofluids were assessed, two different sets of machine learning models were built for both the plasma and saliva samples. The different disease states (HNS, HS, DS, and DA) were assigned different outcome states (0, 1, 2, and 3, respectively) as the outputs of the classifier models. Tables 3 and 4 highlight the classifiers available in the Python library employed to distinguish between the different output states based on the inflammatory markers available for plasma and saliva, respectively.

**Table 3.**
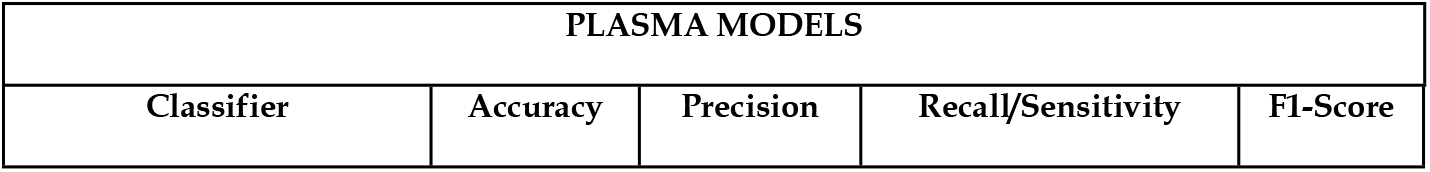
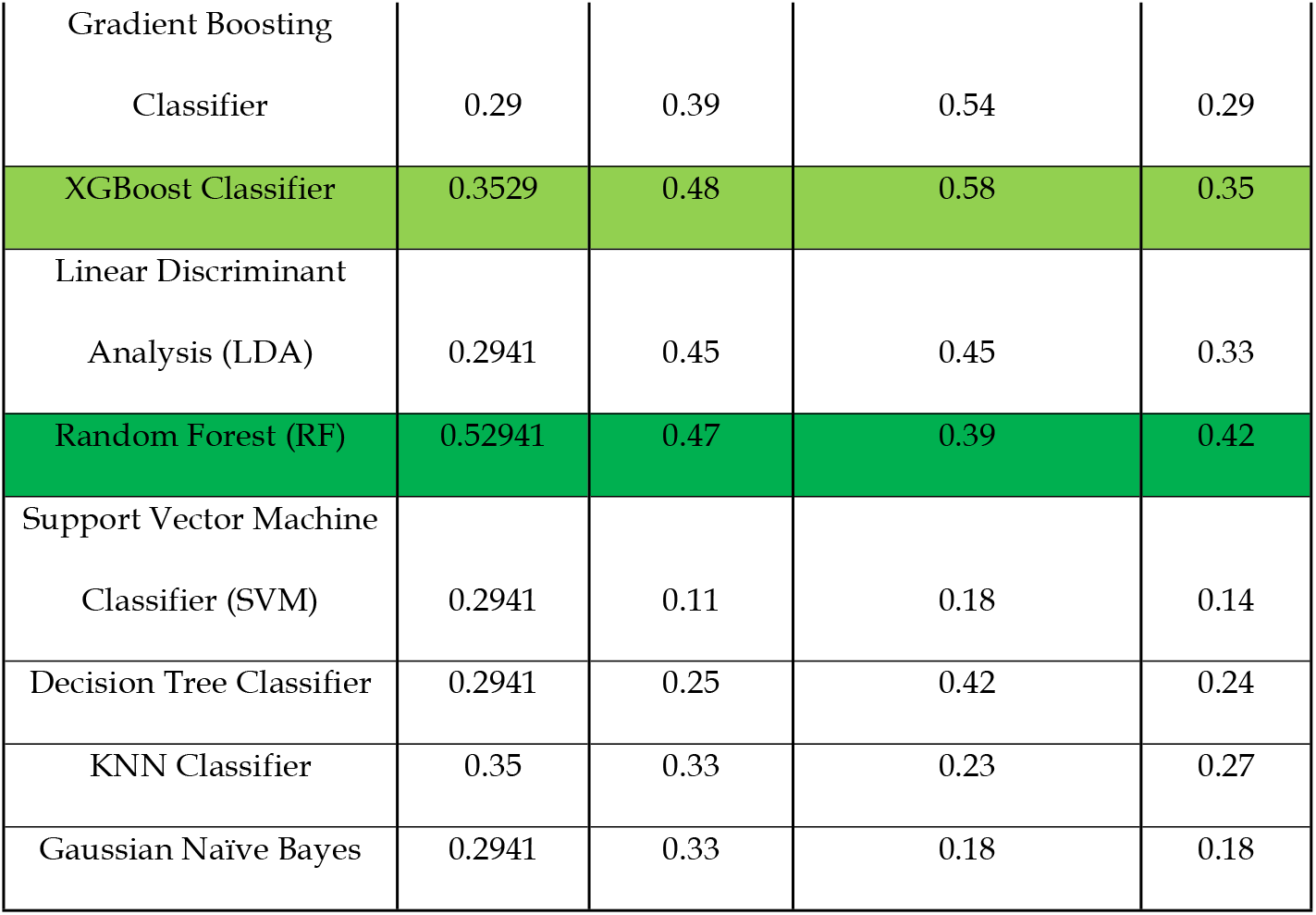
Performance characteristics ML models for classification using biomarkers in plasma.

**Table 4.** Performance characteristics ML models for classification using biomarkers in saliva.

| SALIVA MODELS |  |  |  |  |
| --- | --- | --- | --- | --- |
| Classifier | Accuracy | Precision | Recall/Sensitivity | F1-Score |
| Gradient Boosting Classifier | 0.5 | 0.5 | 0.58 | 0.52 |
| XGBoost Classifier | 0.25 | 0.19 | 0.21 | 0.2 |
| Linear Discriminant Analysis<br>(LDA) | 0.38 | 0.33 | 0.46 | 0.38 |
| Random Forest (RF) | 0.625 | 0.5 | 0.54 | 0.52 |
| Support Vector Classifier (SVC) | 0.625 | 0.75 | 0.71 | 0.67 |
| Decision Tree Classifier | 0.38 | 0.33 | 0.33 | 0.33 |
| KNN Classifier | 0.75 | 0.79 | 0.79 | 0.78 |
| Gaussian Naïve Bayes | 0.375 | 0.18 | 0.29 | 0.23 |

Despite the limited plasma data, we still wanted to identify if any non-linear relationships existed among the different groups. To evaluate the models’ effectiveness in distinguishing between these groups, we assessed their performance by considering, in order of increasing importance: accuracy (the model’s correctness in predicting the right output state based on the measured inflammatory markers), precision (the frequency with which the classifier correctly identifies the groups), recall/sensitivity (the proportion of actual correct predictions relative to all predictions), and F1-Score (which reflects the balance between precision and recall).

As anticipated, the classifier models performed poorly in the plasma models, even with optimal parameter adjustments. The Random Forest (RF) model was the most accurate, achieving around 52% accuracy and 47% precision. However, its sensitivity was relatively low at about 39%. The next best plasma model, XGBoost, only reached an accuracy of approximately 35%. Given the limited dataset available with the plasma samples, such low performance was anticipated.

The Random Forest (RF) classifier’s relatively better performance than the others could be attributed to its ability to handle complex interactions between features through the use of multiple decision trees. This ensemble method reduces overfitting by averaging the results of multiple trees, making it more robust to variability in the data(36). Fig 1 shows the confusion matrix visually depicting the correct prediction of the classification of the top two most accurate plasma models. The confusion matrix for the RF classifier shows that it was able to correctly classify a significant number of samples, but there were still misclassifications, particularly in distinguishing between certain classes.

**Fig 1.**
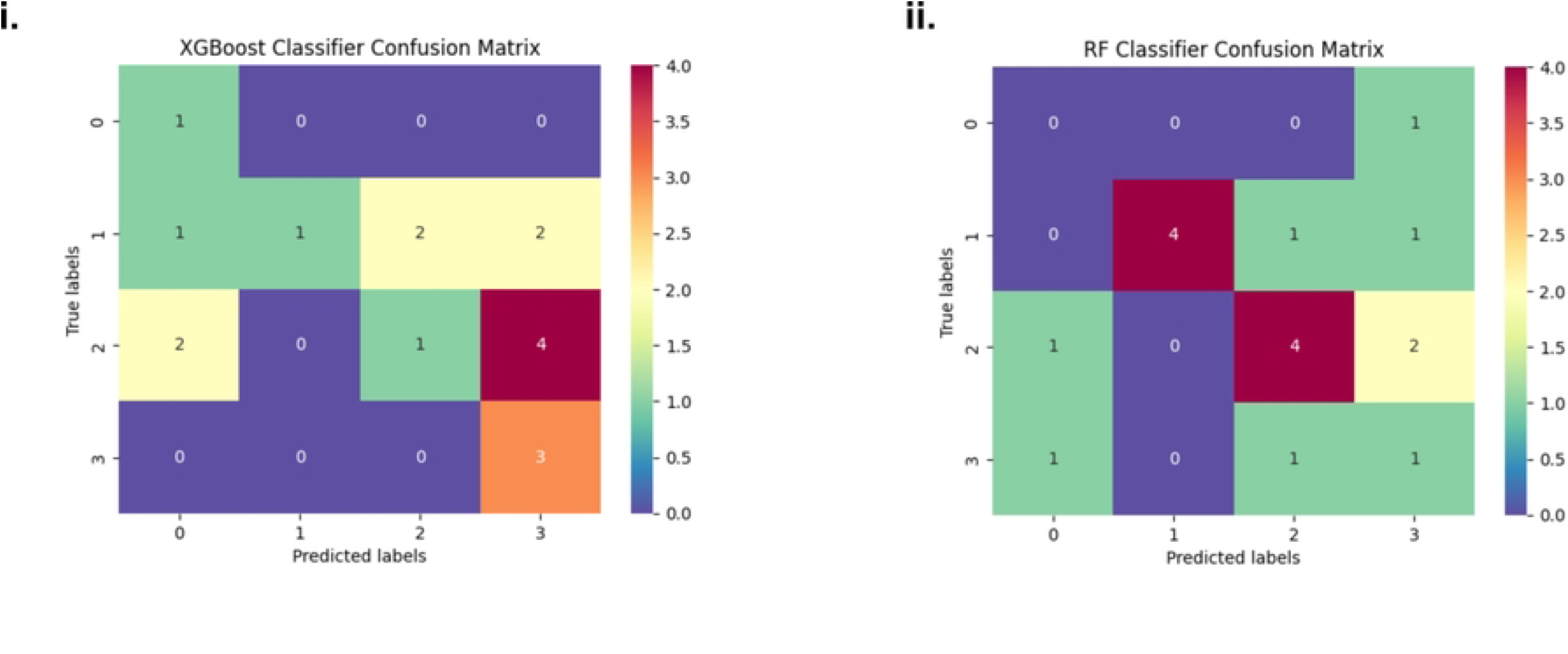
Confusion matrix for the top two best performing models using biomarkers from plasma

We applied the same classifiers to the saliva dataset to evaluate their effectiveness in distinguishing between the four groups, anticipating better classifier metrics due to the increased number of input markers as compared to the plasma data. Using the same metrics as for the plasma dataset, we analyzed the results presented in Table 4. The K-Nearest Neighbor (KNN) Classifier achieved the highest performance, with an accuracy of 75%, a precision of 79%, and a sensitivity of 79%, leading to an F1-Score of 78%. This performance significantly surpassed that of the next best classifier, the Support Vector Machine (SVM) Classifier which had an accuracy of 62.5%. While the random forest model for the saliva model had a better accuracy than the plasma model, it was not the best performing classifier. The K-Nearest Neighbors (KNN) algorithm excels in performance due to its ability to capture local patterns by considering the nearest neighbors(37). The algorithm determines the label of a data point by evaluating nearby data points, making it highly effective for datasets with distinct clusters and since the saliva dataset had more neighbors to consider, it wasn’t surprising to see it outperform the other classifiers. The confusion matrixes for the top two saliva models are shown in Fig 2. below. Comparing the two, we see that the KNN classifier correctly classified most of the samples, indicating the efficacy of the classifier in capturing the local patterns in the saliva data as evidenced by the relatively higher accuracy and precision.

**Fig 2.**
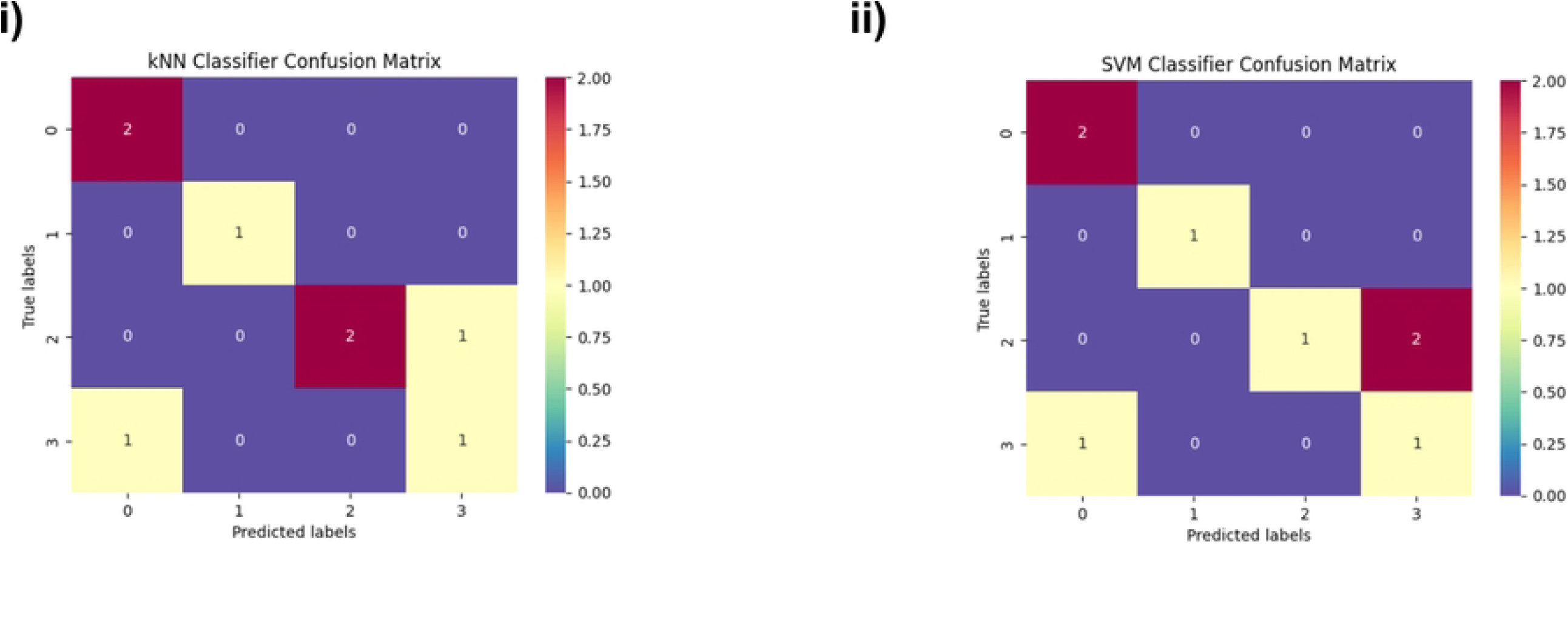
Confusion matrix for the top two best performing models using biomarkers from plasma

To enhance the efficacy of the models, we employed feature selection using the MRMR (Minimum Redundancy Maximum Relevance) ranking. We rerun the models with an increasing number of markers based on the MRMR ranking for both the plasma and saliva models (see Supplementary Information). As expected, there was no significant change in the classifier metrics for the plasma models with the addition of more markers according to the MRMR ranking. However, for the saliva models, using accuracy as the metric, we observed that the accuracy of the best-performing saliva model (KNN) increased with the addition of markers from the MRMR ranking, plateauing at 75% with the first 12 markers as seen in Fig 3. below.

**Fig 3.**
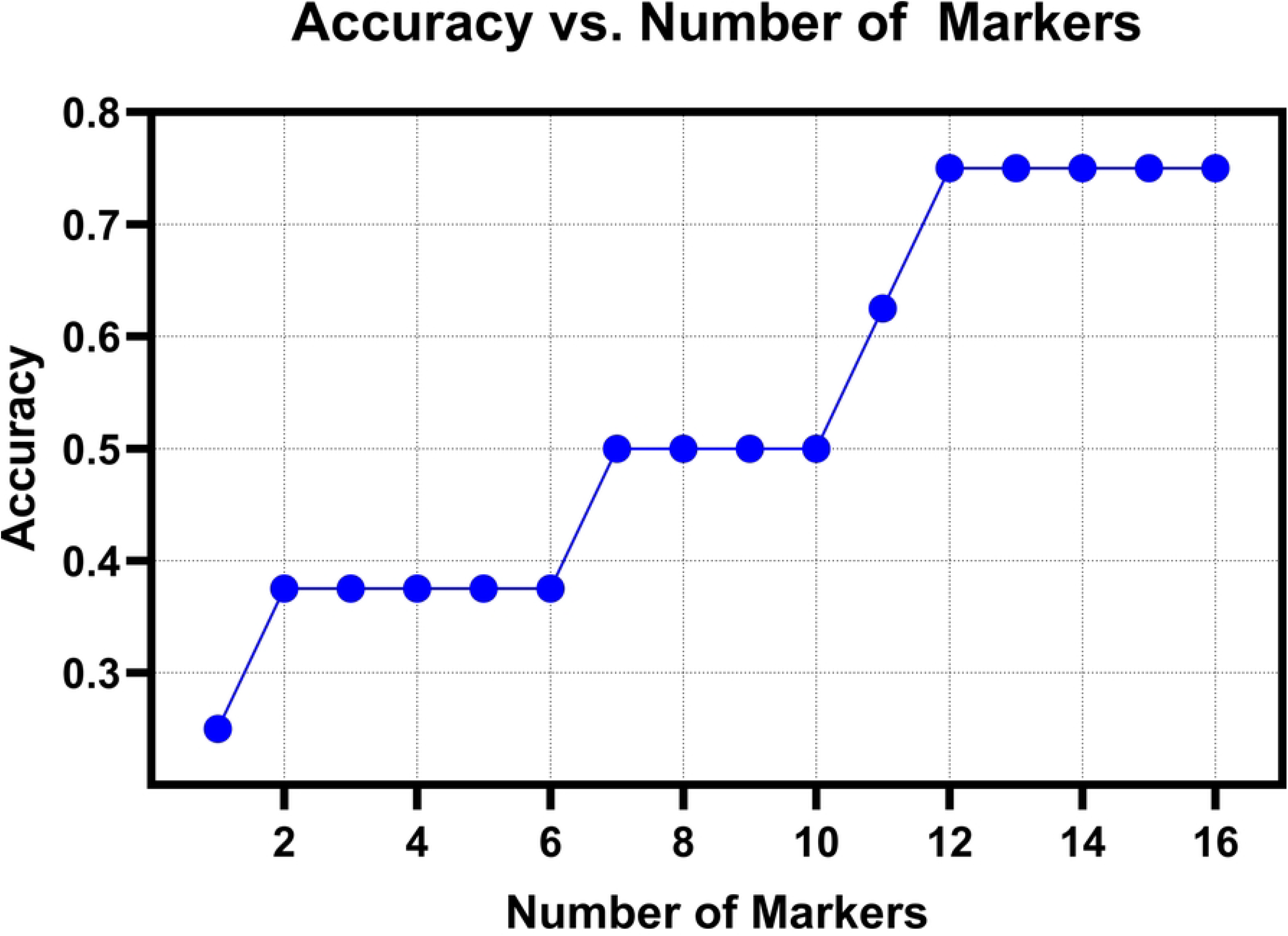
Accuracy plot showing the change in accuracy with increasing number of markers added to the kNN model

Comparing the original model with all 17 markers to the model with the first 12 markers from the MRMR ranking (see Table 5), we observed improved performance metrics in terms of precision and sensitivity. This suggests that using models to identify key markers can enhance the development of personalized medicine strategies.

**Table 5.**
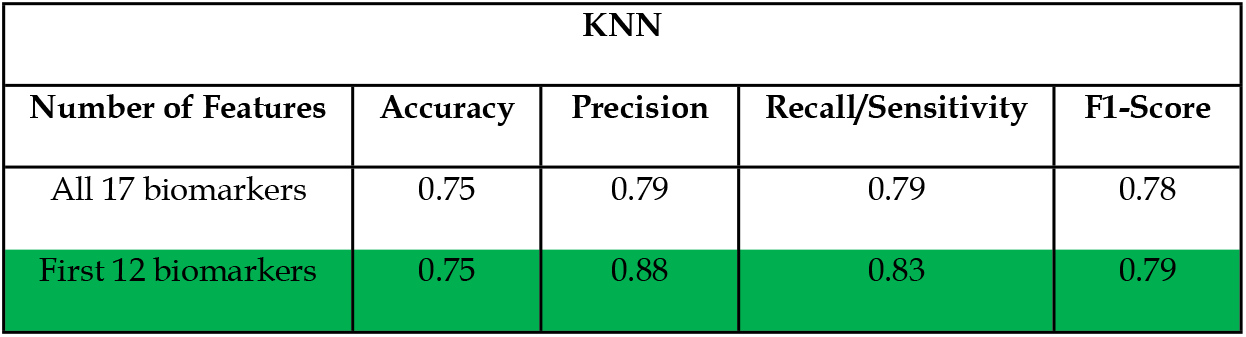
Performance characteristics ML models for classification using biomarkers in saliva with modified number of features.

| KNN |  |  |  |  |
| --- | --- | --- | --- | --- |
| Number of Features | Accuracy | Precision | Recall/Sensitivity | F1-Score |
| All 17 biomarkers | 0.75 | 0.79 | 0.79 | 0.78 |
| First 12 biomarkers | 0.75 | 0.88 | 0.83 | 0.79 |

Machine Learning (ML) has significantly enhanced the classification and management of asthma by providing advanced tools to identify risk factors, predict exacerbations, and optimize treatment strategies. Both Jayamini et al. and Molfino et al. have conducted extensive systematic reviews of the literature on the use of ML algorithms to predict asthma exacerbations, contributing to better control and management of asthma(38,39) where they detail the different ML algorithms that have been employed in asthma modeling, the datasets used and their resulting outcomes. These studies typically rely on large datasets, particularly Electronic Health Records (EHR), which include demographic data, lab results, medication information, and records of exacerbations. The results from these studies demonstrate varying degrees of success in predicting asthma-related events, depending on the size and quality of the dataset used(40,41). Many studies have successfully utilized electronic health records (EHRs), demonstrating their potential in developing robust predictive models. However, to the best of our knowledge, this paper represents one of the first efforts to explore the exclusive use of cytokine expression in plasma and saliva as a sole method for classifying different asthmatic states. While our study’s results are still preliminary, with a limited sample size, the comparative outcomes obtained from our models show promising potential. These early findings suggest that non-invasive saliva cytokine measurements could serve as an effective tool for asthma classification, providing a new avenue for diagnostic and prognostic screening.

## Conclusion

As with all machine learning models, having more data to train and tune them can significantly enhance their effectiveness and efficiency in identifying non-linear relationships between markers. In the context of phenotyping and personalizing treatment for different asthmatic conditions, the identification of key markers is crucial. This study demonstrates the potential of machine learning methods to identify markers that can discriminate between different disease states in asthma. Our findings highlight that saliva inflammatory markers provide a better indication of airway inflammation compared to plasma inflammatory markers, which tend to reflect a more systemic response. This insight is particularly valuable for developing customizable point-of-care devices that can test personalized marker selection based on machine learning models. Such devices would not only validate the models but also provide additional data to further train these models, ultimately improving their accuracy and reliability. By leveraging machine learning to identify critical markers, we can enhance the personalization of asthma treatment and management, leading to better patient outcomes and more targeted therapeutic strategies. This approach holds promise for advancing our understanding of asthma pathophysiology and developing innovative solutions for its treatment and management.

## Acknowledgments

I would like to acknowledge the Eugene McDermott Graduate Fellowship for their help and support in the collection of samples from Ghana. Special thanks to the team at both KNUST and KCCR that aided in the day-to-day collection of the samples.

